# Cryo-Electron Microscopy Structure of the *Macrobrachium rosenbergii* Nodavirus Capsid at 7 Angstroms Resolution

**DOI:** 10.1101/103424

**Authors:** Kok Lian Ho, Chare Li Kueh, Poay Ling Beh, Wen Siang Tan, David Bhella

**Affiliations:** Department of Pathology, Faculty of Medicine and Health Sciences, Universiti Putra Malaysia, 43400 UPM Serdang, Selangor, Malaysia.; Department of Microbiology, Faculty of Biotechnology and Biomolecular Sciences, 43400 UPM Serdang, Selangor, Malaysia.; MRC-University of Glasgow Centre for Virus Research, Sir Michael Stoker Building, Garscube Campus, 464 Bearsden Road, Glasgow G61 1QH, Scotland, UK.

**Keywords:** *Macrobrachium rosenbergii* nodavirus, capsid, cryo-electron microscopy, virus-like particles, 3-dimensional structure

## Abstract

White tail disease in the giant freshwater prawn *Macrobrachium rosenbergii* causes significant economic losses in shrimp farms and hatcheries and poses a threat to food-security in many developing countries. Outbreaks of *Macrobrachium rosenbergii* nodavirus (*Mr*NV), the causative agent of white tail disease (WTD) are associated with up to 100% mortality rates. Recombinant expression of the capsid protein of *Mr*NV in insect cells leads to the production of VLPs closely resembling the native virus. We have investigated the structure of *Mr*NV VLPs by cryogenic electron microscopy, determining a structure of the viral capsid at 7 angstroms resolution. Our data show that *Mr*NV VLPs package nucleic acids in a manner reminiscent of other known nodavirus structures. The structure of the capsid however shows striking differences from insect and fish infecting nodaviruses, which have been shown to assemble trimer-clustered T=3 icosahedral virus particles. *Mr*NV particles have pronounced *dimeric* blade-shaped spikes extending up to 6 nm from the outer surface of the capsid shell. Our structural analysis supports the assertion that *Mr*NV along with the related virus of marine shrimp *Penaeus vannamei* nodavirus (*Pv*NV) may represent a new genus of the *Nodaviridae.*

**Author summary:** *Macrobrachium rosenbergii* nodavirus (*Mr*NV) is the causative agent of white tail disease (WTD) which leads to 100% mortality in shrimp-farms growing giant freshwater prawn (*M. rosenbergii*). *Mr*NV is therefore a significant threat to food security and causes severe economic losses in developing countries such as Malaysia, Indonesia, Pakistan, Thailand and India. Here we have used electron microscopy to study the three-dimensional structure of *Mr*NV, revealing that the viral capsid – the protein shell that encloses the viral genome, protecting it and transporting it from one host to the next – is differently organised to capsids produced by other viruses in the nodavirus family. The virus was found to have large blade-like spikes on its outer surface that are likely important in the early stages of infection, when the virus attaches to and enters a host cell.

## Introduction

*Macrobrachium rosenbergii* nodavirus (*Mr*NV) causes white tail disease (WTD) in the giant freshwater prawn *M. rosenbergii. Mr*NV outbreaks in shrimp farms and hatcheries can result in mortality rates of up to 100%, thus the virus represents a threat to food-security and causes significant economic losses [1]. WTD disease was first reported in Pointe Noire, Guadeloupe in 1997, and was subsequently detected in China [2], India [3] Taiwan [4], Thailand [5], Malaysia [6], Australia [7] and most recently in Indonesia [8].

*Mr*NV belongs to the family *Nodaviridae*, which is divided into *Alphanodavirus* and *Betanodavirus* genera. The former includes insect-infecting nodaviruses such as Nodamura virus (NoV), Boolaara virus (BoV), Flock House virus (FHV), black beetle virus (BBV) and Pariacoto virus (PaV). The latter comprises fish-infecting nodaviruses such as Malabaricus grouper nervous necrosis virus (MGNNV) [see [6] for a review]. Based on the amino acid sequence of the RNA dependent RNA polymerase (RdRp), *Mr*NV is grouped within the *Alphanodavirus* genus but owing to some unique characteristics of *Mr*NV, a new genus (*Gammanodavirus*) has been proposed which would comprise *Mr*NV and *Penaeus vannamei* nodavirus (*Pv*NV) [9]. In common with other nodaviruses, *Mr*NV is a non-enveloped, icosahedral virus which comprises a nucleocapsid containing a bipartite positive-sense RNA genome. The genomic RNAs: RNA 1 (3.2 kb) and RNA 2 (1.2 kb), encode the RdRp and capsid protein, respectively. The full length *Mr*NV capsid protein is a polypeptide containing 371 amino acids and having an N-terminal region rich in positively charged amino acids that is believed to be an RNA binding region [10]. The C-terminal region of the capsid protein is involved in host cell binding and internalization [11]. This feature allows foreign epitopes such as the ‘*a*’ determinant of hepatitis B surface antigen (HBsAg) and the ectodomain of influenza A virus M2 to be inserted at the C-terminal region and thereby displayed on the surface of the *Mr*NV capsid [12, 13]. Other functional regions within the polypeptide of *Mr*NV capsid protein have not been clearly identified.

Recombinant *Mr*NV capsid protein produced in *Escherichia coli* and *Spodoptera frugiperda* (both *Sf*9 and *Sf*21) assembles into icosahedral virus-like particles (VLPs) although these were found to be morphologically different under transmission electron microscopy analysis [14, 15]. Negative stained VLPs expressed in *E. coli* and *Sf*9 were measured to have diameters of ~30 and 40 nm, respectively [14, 15]. The self-assembly/disassembly capability of the *Mr*NV capsid protein has been exploited to develop nano-carriers for DNA and double stranded RNA [16, 17]. Owing to the economic importance of *Mr*NV and the potential applications of recombinant capsid proteins, there is considerable interest in the three-dimensional (3D) structure of the *Mr*NV nucleocapsid. Determining the structure will provide insights into the molecular details of virion morphogenesis, potentially leading to the identification of *Mr*NV inhibitors suitable for use in combating *Mr*NV infection. To date, the high-resolution structures of several alphanodaviruses including FHV [18], PaV [19], BBV [20] and NoV [21] have been solved by X-ray crystallography. Of the betanodaviruses, a high-resolution structure has been determined for grouper nervous necrosis virus (GNNV) [22]. A structure for the unclassified Orsay virus, which infects the nematode worm *Caenorhabditis elegans*, has also been determined [23]. All known structures of *Nodaviridae* reveal T=3 icosahedral capsids assembled from 180 capsid proteins that have an eight-stranded antiparallel beta-barrel topology. Known nodavirus capsid structures display trimer-clustering; PaV and NoV having small trimeric spikes on their outer surface, while Orsay and GNNV have larger trimeric capsomeres.

Amino acid sequence alignment showed that FHV capsid protein shares ~87% sequence identity with BBV capsid protein. However, none of these nodavirus capsid proteins show similarities beyond 20% with *Mr*NV or *Pv*NV capsid proteins. Here, we report a 3D reconstruction of *Mr*NV VLPs determined by cryo-electron microscopy at a resolution of 7 Å. The 3D reconstruction reveals a T=3 icosahedral assembly that shows a striking divergence from the known structures of other nodavirus capsids. This was characterized by the presence of large dimeric blade-like spikes on the outer surface. Although the VLPs assembled in a heterologous system and in the absence of the full-length viral genome, we find that the capsids contain density consistent with the encapsidation of RNA, which we assume to be the cognate mRNA. Overall, the considerable morphological differences seen when comparing the structure of *Mr*NV to other known nodaviruses supports the assertion that *Mr*NV and *Pv*NV, which share ~45% sequence similarity in their capsid proteins, represent a distinct genus of nodaviruses.

## Results

### *Mr*NV VLPs assemble as dimer-clustered T=3 icosahedral capsids that package nucleic acids

To investigate the structure of the *Mr*NV capsid, VLPs were produced by expression of the capsid gene in a recombinant baculovirus system and purified from *Sf*9 cells by differential ultracentrifugation. VLPs were prepared for cryo-electron microscopy by plunge freezing in liquid-ethane. Figure 1(a) shows a representative cryomicrograph revealing a population of particles with pronounced spikes on their outer surface and that measured 40 nm in diameter. VLPs appeared dense in projection, suggesting that they contain nucleic acids. Two hundred and eighty-one cryo-micrographs of *Mr*NV VLPs were recorded for 3D image reconstruction. Automated particle picking was used to extract a dataset of 9592 particles. These data were processed to calculate a reconstruction using the RELION software package. Following 2D and 3D classification procedures, 1658 particles were retained for the final round of refinement, achieving a resolution of 7 angstroms (Supplemental Figure 1).

**Figure 1.**
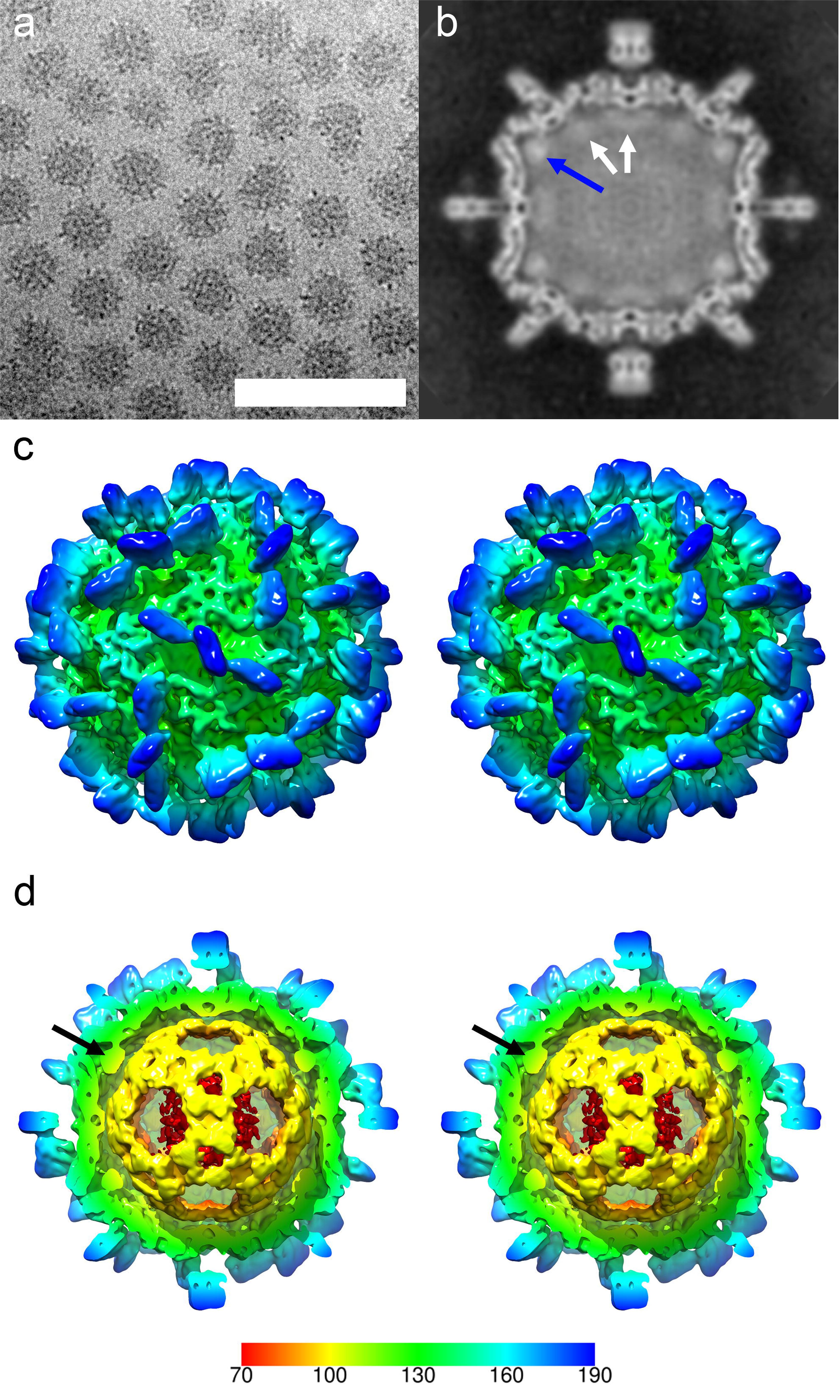
(a) Cryo-EM of *Mr*NV VLPs revealed a population of isometric particles with pronounced spikes on the outer surface. VLPs were dense in projection, indicating that they packaged nucleic acids (Scale bar 100 nm). (b) A central section through the reconstructed *Mr*NV VLP density map shows a sharply defined capsid structure with pronounced spikes on the outer surface. Lining the capsid interior is a region of fuzzy density (white arrows) that we attribute to packaged RNA. The strongest density is seen to lay under the icosahedral five-fold symmetry axes (blue arrow). (c) A wall-eyed stereo-pair image of the isosurfaced 3D reconstruction highlights the overall particle morphology featuring pronounced dimeric spikes on the capsid surface at the two-fold symmetry axes and arranged about the five-fold axes. (d) A cut-away view reveals that the internal density highlighted in (b) forms a dodecahedral cage, a feature previously documented in other nodavirus structures. The density underlying the five-fold symmetry axes is not, however, contiguous with this structure (black arrow). The colour key indicates the scale (in angstroms) of the radial colour scheme used in figures 1 (c,d), 2 and 3 (a,c).

Figure 1(b) shows a central slice through the reconstruction prior to sharpening and figure 1(c) is a stereo-pair view of the isosurfaced reconstruction, also unsharpened. The density map reveals a particle with pronounced, blade-like, 6 nm spikes on the outer surface that are located at the icosahedral two-fold symmetry axes and about the five-fold axes. The location of these spikes is consistent with dimer-clustered T=3 icosahedral symmetry. Thus, we shall adopt the domain nomenclature used when describing similar structures, such as capsids of the *Caliciviridae*. We divide the capsid protein into protruding (P) domain and shell (S) domain [24]. Likewise, we shall describe the capsid structure as having dimers assembled from three quasi-equivalent capsid proteins termed A, B and C. These are arranged as AB dimers (about the five-fold axes) and CC dimers (located at the two-fold axes).

Inspection of figure 1(b) reveals a sharply resolved capsid shell measuring between 2 and 3.5 nm thick. Lining the capsid interior is a less well resolved fuzzy density (white arrows) that is particularly bright beneath the five-fold symmetry axes (blue arrow). We assume this to be packaged nucleic acids, most likely RNA. In figure 1(d) the reconstruction is cropped to reveal this internal density, showing the presence of a dodecahedral cage structure. Similar features have been described previously in cryo-EM studies of nodaviruses and other positive sense RNA containing viruses [19, 25–28]. The density below the five-fold symmetry axes is not, however, contiguous with this feature (black arrow).

### *Mr*NV P-domains form blade-like dimers

The most striking feature of the *Mr*NV capsid is the presence of large blade-like P-domains. These features are well resolved in the reconstructed density, and show marked differences between spikes at the two-fold axes (CC dimers) and those about the five-fold axes (AB dimers).

Figure 2(a) shows the *Mr*NV VLP reconstruction following a sharpening procedure that reduces the influence of low-resolution information on the isosurface representation, thus enhancing high-resolution detail. A close-up view of the AB and CC dimers is presented in figures 2(b,c) and 2(d) respectively. These images show that the P-domain dimers form square-shaped blades that are 4.7 nm in both width and height. The spikes are however very thin, measuring only 1.8 nm in depth. These structures are clearly two-fold symmetric as can be seen in figures 2(b) and (c), in which the AB spike is rotated 180° (unlike the CC spike, rotational symmetry is not applied to this feature by the icosahedral reconstruction process as it is located at a quasi-two-fold symmetry axis). The density of the spikes shows comparable features in both AB and CC dimers except that the P-domains are quite differently oriented relative to their respective S-domains. Firstly, the CC P-dimers are raised from the capsid surface on two well resolved legs of density and with a clearance of approximately 2 nm from the outer surface of the capsid shell. The AB P-dimers sit much closer to the outer surface of the capsid shell and are tilted approximately 20° towards the icosahedral two-fold symmetry axes such that on the two-fold proximal side of the feature, there is ~0.5 nm between the P-domains and the S-domains, while on the 2-fold distal side, a gap of ~1.5 nm separates them. At this resolution, the AB P-dimers appear to contact their nearest CC P-dimers (black arrow figure 2b), creating a larger structure comprised of three P-dimers (two AB P-dimers and one CC P-dimer).

**Figure 2.**
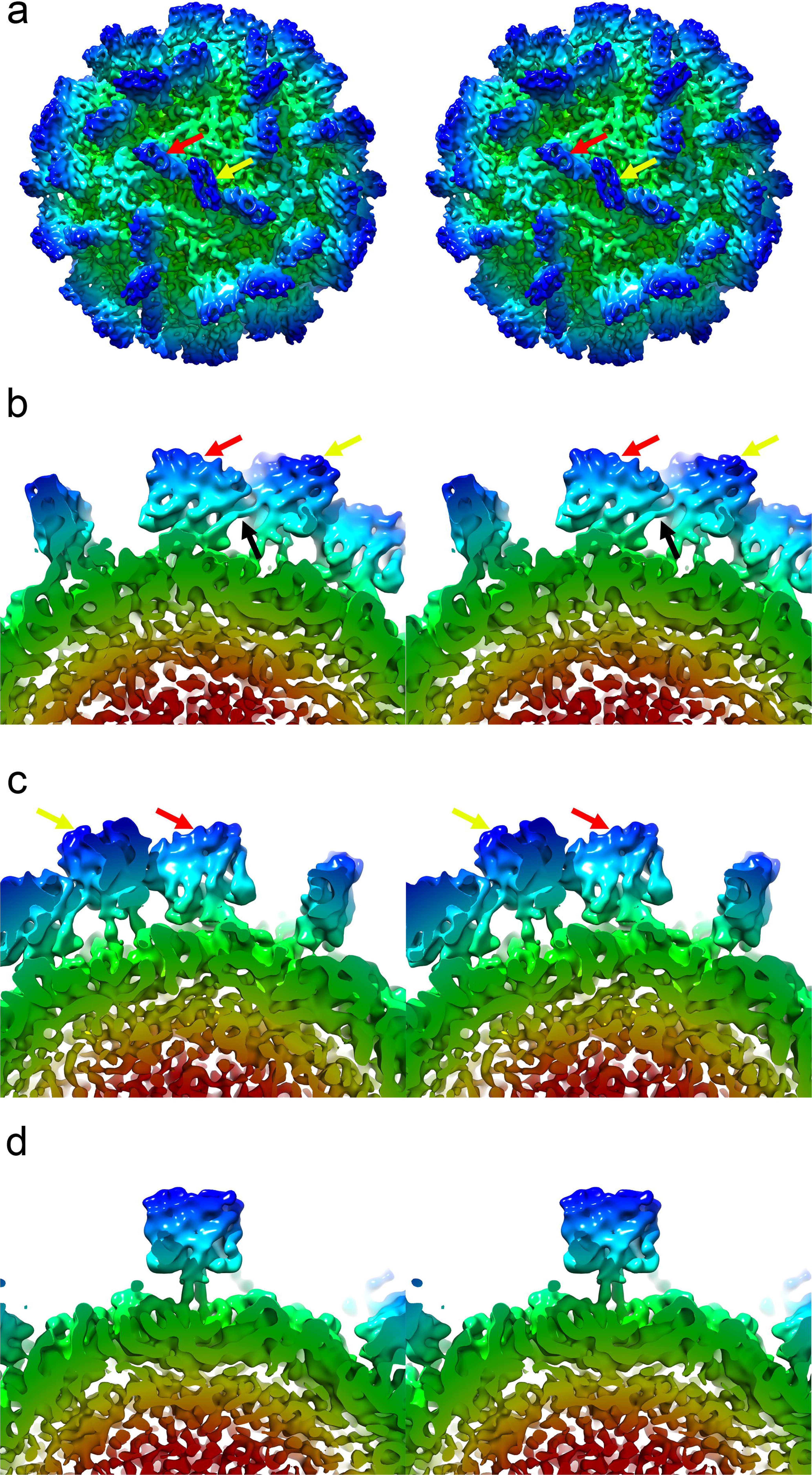
(a) Stereo view of the sharpened 3D reconstruction of the *Mr*NV VLP showing the location of AB (red arrow) and CC (yellow arrow) P-dimers. (b) An AB P-dimer (red arrow), viewed parallel to the capsid surface, showing that the protruding spike leans towards the icosahedral two-fold symmetry axis, making contact with the CC-dimer, which is identified by a yellow arrow, the point of contact is highlighted with a black arrow. (c) A view of the same AB P-dimer shown in (b), but rotated by 180°, highlighting the two-fold symmetry of this feature – a property that is not imposed by the 3D reconstruction process. (d) unlike the AB-dimer, the CC dimer is oriented perpendicular to the capsid surface and raised on two clearly resolved legs of density.

Another striking feature of the P-domains is that there is a pronounced rotation in the orientations of the AB and CC P-dimers relative to the underlying S-domains and the icosahedral symmetry axes. Figure 3(a) shows a stereo-pair image of the *Mr*NV VLP viewed along an icosahedral three-fold symmetry axis. In T=3 icosadeltahedra, quasi-six-fold symmetry axes are located at the icosahedral three-fold axes, as highlighted by the ‘soccer ball’ cage of pentagons and hexagons indicating the locations of icosahedral five-fold and local six-fold symmetry axes. We therefore expect gross morphological features, such as domain orientation to follow this local six-fold symmetry as is seen in the dimer-clustered T=3 capsids encoded by the *Caliciviridae*. Figure 3(b) shows a structure for feline calicivirus (FCV) that we have recently determined [29]. In this reconstruction, the orientation of the CC P-dimer (orange arrow) is approximately related to that of the AB P-dimer (yellow arrow) by a 60° rotation about the local six-fold symmetry axis (indicated by a red hexagon). Figure 3(c) highlights the relative orientations of the P-domains for *Mr*NV, showing that the orientations of the CC P-dimers deviate by approximately 105° relative to that of the AB P-dimers, such that the orange arrows indicating the orientations of the CC P-dimers point towards the local six-fold axis, while the yellow arrows, marking the orientations of the AB P-dimers, point clockwise relative to this axis.

**Figure 3.**
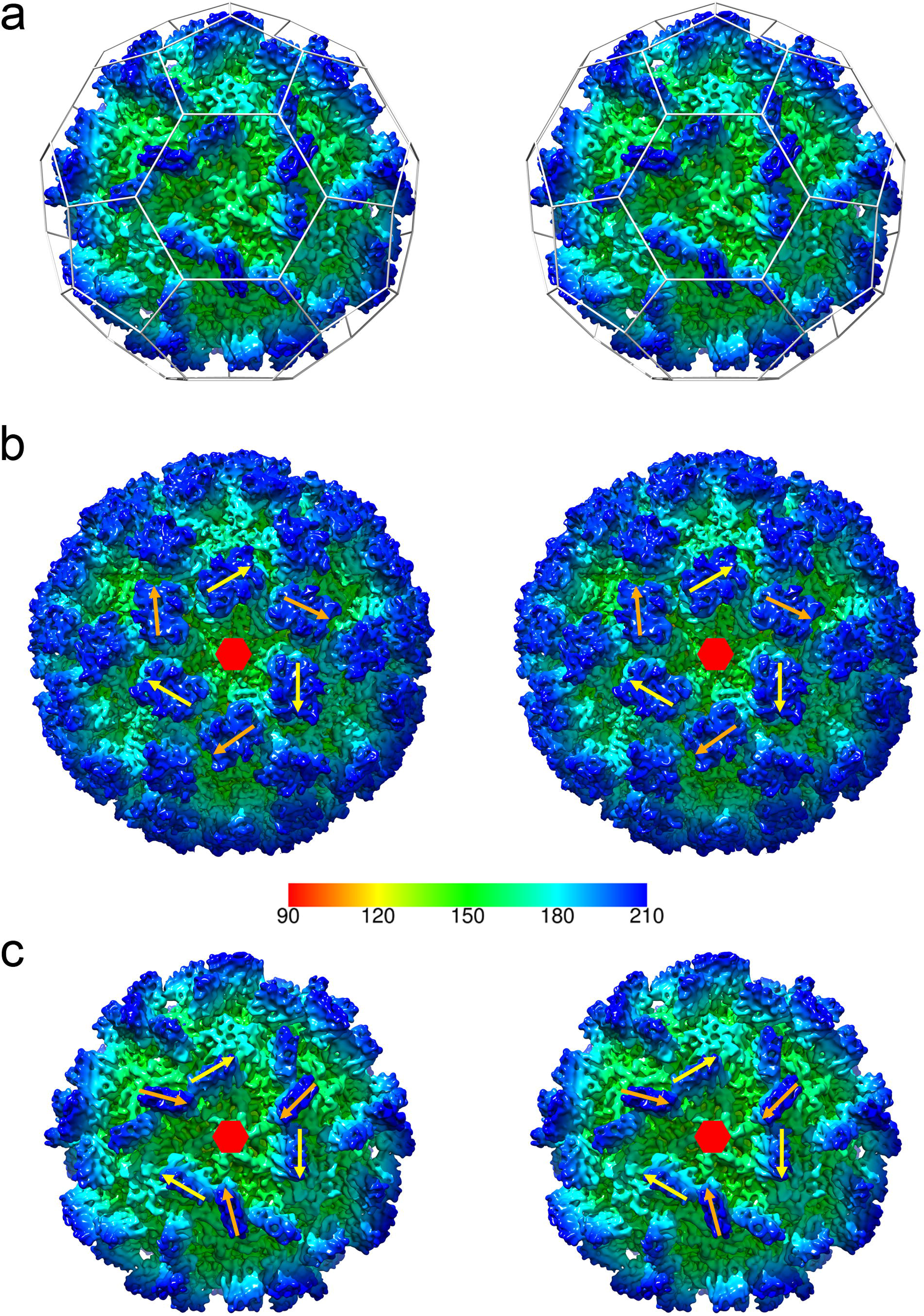
(a) A stereo pair of the *Mr*NV reconstruction viewed along an icosahedral three-fold symmetry axis. In T=3 icosahedral capsids, a local six-fold quasi-symmetry axis lies at this point. The locations of icosahedral five-fold and local quasi-six-fold symmetry axes are highlighted by a ‘soccer-ball’ cage. (b) Dimer clustered T=3 icosahedral capsids, such as our recently published structure for feline calicivirus have AB and CC P-dimers arranged alternating about the quasi-six-fold axes. Here the positions of such are indicated by yellow arrows (AB) and orange arrows (CC). If the structures of AB and CC dimers are not radically different, we expect the position and orientation of these features to be related by an approximately 60° rotation about the local six-fold symmetry axis (indicated by a red hexagon). (c) In the case of the *Mr*NV P-dimers, the CC P-dimers are rotated approximately 105° relative to the orientation of the AB P-dimers. The radial colour scheme in panel (b) indicates the radial depth cue scale in angstroms for the FCV reconstruction only.

## Discussion

*Macrobrachium rosenbergii* nodavirus is an important pathogen that poses both an economic and a food-security threat to the growing fresh-water aquaculture industries of many developing countries. We sought to generate structure data that would support the development of interventions to treat or prevent outbreaks of *Mr*NV disease. Such incidents are associated with mortality rates of up to 100% owing to the ease with which waterborne pathogens can spread among the dense populations found in shrimp farms.

We used cryogenic transmission electron microscopy to image purified *Mr*NV VLPs in a frozen hydrated state. Processing the resulting images led to the calculation of a 7 Å resolution 3D reconstruction of the capsid.

We found that expression of the recombinant capsid protein in insect cells led to the production of VLPs that when reconstructed showed density reminiscent of packaged viral genomes in known nodavirus structures [19, 25]. We hypothesize that this density is packaged nucleic acid, possibly the cognate mRNA. During authentic infections, positive-sense RNA containing viruses do not express their genetic content via the production of mRNA, rather their genomes are directly translated by host ribosomes to produce the encoded gene-products. There is growing evidence that capsid assembly in many positive-sense RNA viruses is directed by specific encapsidation signals throughout the viral genomic RNA [30]. It is possible then that such encapsidation signals may also be present in mRNA produced in experiments such as ours, in which a DNA sequence was generated from the viral genome for recombinant expression.

Although our reconstruction is insufficient to determine the tertiary structure of the capsid protein, it has revealed a striking divergence from known nodavirus structures. Most notably, while other known nodavirus virions exhibit *trimer*-clustering of capsid proteins arranged with T=3 icosahedral symmetry [19, 22, 25, 26], our map shows pronounced *dimeric* capsomeres. Our data further revealed a capsomere morphology that is very different from known similar capsids. Firstly the spikes we observed were seen to form as approximately square, thin blade-shaped protrusions. Moreover there was considerable difference in the orientation of these capsomeres relative to each other. The AB P-domains were seen to lean towards the icosahedral two-fold symmetry axes, while the CC P-domains were raised away from the capsid shell. Furthermore the lean resulted in the AB P-dimers contacting the CC P-dimers to form a larger blade-like super-structure lying across the icosahedral two-fold axes. There was also a rotation of 105° in the quasi-symmetry related orientations of AB and CC P-domains, possibly a consequence of the contact between the two dimer forms.

It has previously been reported that the C-terminal domain of the *Mr*NV capsid protein likely encodes the functional attachment and entry requirements of the virus [11], while the N-terminal region contains positive charged residues that bind the viral genome [10]. Our structure shows that the CC P-dimer is supported on two narrow legs of density. Taken together these data strongly suggest that the P-domain is not an insert in the capsid polypeptide chain rather it is at the C-terminus. The presence of a discrete dimeric P-domain comprising the C-terminal region of the capsid protein was proposed by Somrit *et al.* [11]. In that study web-based homology modelling software was used to predict the structure of the *Mr*NV capsid protein. Protein sequence similarity analysis identified the capsid protein of a tombusvirus - cucumber necrosis virus as a suitable template. A model based on the known crystal structure of that protein [31] was calculated, producing a predicted two-domain structure comprising antiparallel beta-barrels in both S and P-domains (Supplemental Figure 2). This model is superficially compatible with our cryo-EM density map, however when we repeated this analysis (using the Phyre2 server [32]) and attempted to dock the resulting coordinates into our reconstruction using rigid body fitting of individual domains in UCSF Chimera [33], we were unable to achieve a satisfactory fit. Moreover, attempts to use a similar approach with SWISS-MODEL [34] identified capsid proteins of other nodaviruses (Orsay virus [23]) or tombusviruses (carnation mottle virus [35]) as suitable templates for modelling. Once again, attempts to dock the resulting coordinates into our density map did not yield convincing results. While these modelling experiments strongly point to a likelihood of the *Mr*NV capsid protein S-domain adopting an eight-stranded antiparallel beta-barrel topology, they have not allowed us to propose a model that is consistent with our intermediate resolution cryo-EM data.

In summary, our analysis of the structure of the *Mr*NV capsid has demonstrated that this virus (and likely the related *Pv*NV) is highly diverged from the alpha- and beta-nodaviruses, as the capsid shows a unique morphology characterized by the presence of thin blade-like dimeric P-domains, comprising the C-terminus of the capsid protein. These capsomeres come together to form a large linear super-structure on the capsid surface lying across the icosahedral two-fold symmetry axes. This feature is likely critical for virus attachment and entry. These data strongly support the assertion that both *Mr*NV and *Pv*NV may represent a new genus of the virus family *Nodaviridae.*

## Methods and Materials

### Preparation of baculovirus stock

P1 stock of baculovirus was prepared according to the manufacturer’s protocol (Bac-to-Bac Baculovirus Expression System, Invitrogen) with some modifications. Briefly, the P1 stock was prepared by transfecting the *Spodoptera frugiperda* (*Sf*9) cells at a cell density of 8 × 10^5^ cells/well on a 6-well plate with Bacmid containing the *Mr*NV capsid protein gene and a 6× His-tag coding sequence. Following incubation at 27 °C for 72 hours, the medium containing the P1 baculovirus was collected. The remaining insect cells and large debris in the medium were removed by centrifugation at 500 × g for 5 min at 4 °C. The P1 stock of baculovirus was stored at 4 °C, protected from light. To prepare the P2 stock of baculovirus, *Sf*9 cells at a cell density of 2 × 10^6^ cells/well in a 6-well plate were infected with the P1 stock. The infected cells were incubated at 27 °C for 48 hours before the medium containing the P2 baculovirus stock was collected. The remaining insect cells and large debris in the medium were removed by centrifugation at 500 × g for 5 min at 4 °C. The P2 stock of baculovirus was stored at 4 °C before used.

### Expression and purification of His-tagged MrNV capsid protein

*Sf*9 cells were grown at 27 °C in Sf-900 III SFM (Life Technologies, USA) supplemented with 4% (v/v) foetal bovine serum (FBS). When the cell density reached 1–2 × 10^6^ cells/ml, the cells were infected by 10% (v/v) of the recombinant baculovirus stock harbouring the *Mr*NV capsid protein gene and a 6×His-tag coding sequence. The culture was incubated for 4 days before the infected cells and the culture medium were separated by centrifugation at 500 × g for 5 min. The proteins in the culture medium were precipitated at 50% (w/v) ammonium sulfate saturation for 2 hours and the precipitated proteins were pelleted down at 12,000 × g for 20 min and resuspended in sodium phosphate buffer (77.4 mM Na_2_HPO_4_, 22.6 mM NaH_2_PO_4_; pH 7.4). The proteins were then separated by sucrose gradient ultracentrifugation [10 – 40% (w/v)] at 150,000 × g for 4.5 hours at 4 °C with a SW 40 Ti rotor (Beckman Coulter, USA). This was followed by fractionation into 1 ml fractions. The fractions which contained the *Mr*NV capsid protein, as analysed by sodium dodecyl sulphate-polyacrylamide gel electrophoresis (SDS-PAGE), were dialysed in sodium phosphate buffer as above. The purified protein was concentrated by centrifugation using a centrifugal concentrator with molecular cut-off 50 kDa (Microsep centrifugal devices, Pall, USA) at 4000 × g at 4 °C. The final protein concentration was determined by the Bradford assay.

### Cryo-electron microscopy

Purified *Mr*NV capsid protein at approximately 1 mg/ml was prepared for cryogenic transmission electron microscopy using an FEI Vitrobot Mk IV. Four μl of VLP preparation was applied to freshly glow-discharged quantifoil holey carbon support films (R2/2; Quantifoil, Jena, Germany), blotted for 4 seconds and plunged into liquid ethane [36]. Vitrified samples were viewed at low-temperature (around −150 K) and under low electron dose conditions using a JEOL 2200 FS cryo-microscope operated at 200 kV. Samples were held in a Gatan 626 cryo-stage. Micrographs were recorded at 50,000 × magnification on a Direct-Electron DE20 DDD camera giving a sample rate of 1.09 Å/pixel. Three-second exposures, at an electron dose rate of ~20 e/Å^2^/s, were captured in movie mode running at 20 frames per second.

### Computational image reconstruction

Micrograph movies containing 60 frames were processed to calculate a 3D reconstruction using Relion-2.0 [37]. Initially movies were corrected for specimen movement using MotionCorr [38] and defocus was estimated using GCTF [39]. An initial set of ~1000 particles was manually picked and processed to calculate a 2D average suitable for automated particle picking. Following automated particle picking 2D classification was used to select intact particles for 3D classification and autorefinement, imposing icosahedral symmetry. For 3D classification, a starting model was prepared from the X-ray crystallographic coordinates of the Flock House virus capsid protein (PDB 4FTB – DOI 10.2210/pdb4ftb/pdb) using the EMAN program pdb2mrc [40]. In line with recommended procedures the starting model was low-pass filtered to 60 Å to limit the effects of model-bias. Following refinement, per-particle movie correction was performed using only frames 2-20, to limit the impact of radiation damage. Finally, a post-processing b-factor of -890 Å^2^ was applied to the reconstruction [41]. The calculated 3D reconstruction was visualized in UCSF chimera and IMOD [33, 42].

## Author contributions

Conceptualization: KLH, WST, DB

Data Curation: KLH, DB

Investigation: KLH, CLK, PLB, DB

Formal analysis: KLH, DB

Methodology: DB

Visualization: DB

Project Administration: KLH

Writing – Original Draft Preparation: KLH, DB

Writing – Review and Editing: KLH, WST, DB

Funding acquisition: KLH, DB

Resources: WST, DB

Supervision: KLH, WST, DB

## Acknowledgement

We thank Dr. Chyan Leong Ng for discussions and critical reading of the manuscript, KLH was sponsored by Universiti Putra Malaysia (UPM). CLK and PLB were supported by the Graduate Research Fellowship, UPM and MyBrain, the Ministry of Higher Education, Malaysia. This work was supported by UPM Putra Grant (Project number: GP-IPS/2016/9509400) and the United Kingdom Medical Research Council (MC_UU_12014/7).

## Figure legends

Figure S1.

Gold-standard Fourier shell correlation plot for the *Mr*NV 3D reconstruction, indicating a resolution of 7 angstroms was achieved assuming an FSC cut-off of 0.143.

Figure S2

Homology modeling of the *Mr*NV capsid protein. Structure prediction was performed using the known structures determined by X-ray crystallography. The left-hand panels show ribbon diagrams of the homology models, while the right-hand panels show solvent excluding surface representations of the viral capsid structures that informed the homology model. (a,b) cucumber necrosis virus – PDB ID 4LLF, (c,d) cocksfoot mottle virus 2 – PDB ID 1NG0, (e,f) Orsay virus – PDB ID 4NWV and (g,h) carnation mottle virus PDB ID 1OPO. These models show a common eight-stranded anti-parallel beta-barrel topology in the S-domain, however attempts to dock these models to our 3D reconstruction yielded low correlation values. The N and C terminal residues for each ribbon diagram are labelled, indicating the extent of the capsid sequence that was modelled.

